# Bilayer graphene oxide membranes for a wearable hemodialyzer

**DOI:** 10.1101/2020.05.26.116863

**Authors:** Richard P. Rode, Henry H. Chung, Hayley N. Miller, Thomas R. Gaborski, Saeed Moghaddam

## Abstract

2D nanomaterials have long been considered for development of ultra-high throughput membranes, due to their atomically thin nature and high mechanical strength. However, current processes have yet to yield a viable membrane for practical applications due to the lack of scalability and substantially improved performance over existing membranes. Herein, a graphene oxide (GO) bilayer membrane with a permeability of 1562 mL/hr.mmHg.m^2^, two orders of magnitude higher than existing nanofiltration membranes, and a tight molecular weight cut-off (MWCO) is presented. To build such a membrane, we have developed a new process involving self-assembly and optimization of GO nanoplatelets physicochemical properties. The process produced a highly organized mosaic of nanoplatelets enabling ultra-high permeability and selectivity with only three layers of GO. Performance of the membrane has been evaluated in a simulated hemodialysis application, where it presents a great value proposition. The membrane has a precise molecular cut-off size of 5 nm, adjusted using a molecular interlinker, designed to prevent loss of critical blood proteins. Urea, cytochrome-c, and albumin are used as representative test molecules. Urea and cytochrome-c sieving coefficients of 0.5 and 0.4 were achieved under physiological pressure conditions, while retaining 99% of albumin. Hemolysis, complement activation, and coagulation studies exhibit a performance on par or superior to the existing hemodialyzer materials.

---

Fabrication of high-throughput and selective separation media at low cost has been the main objective of the membrane industry for decades. Recent advancements in nanomaterials have opened new opportunities for development of such membranes. Graphene and graphene oxide (GO), the pinnacle of ultrathin mechanically strong 2D nanomaterials^1–3^, have long been considered for separation applications. Theoretical studies and simulations have demonstrated that nanoporous graphene and GO exhibit superior sieving potential in water separation applications^4,5^. However, despite their significant promise, their use has not been realized due to the lack of scalable fabrication processes^6^. Development of a facile and scalable process demands new approaches and a holistic understanding of the relation between physicochemical properties of these 2D nanomaterials and the membrane transport characteristics.

A number of fabrication processes have been proposed to produce nanoporous graphene and GO membranes. Porous single layer graphene membranes patterned via ion bombardment^7,8^, focused-ion beam (FIB) irradiation^9^, or focused electron beam (FEB) writing^10,11^ with pore sizes of 0.4-0.6 nm and pore densities in the 10^12^ cm^−2^ range have been fabricated. While these efforts demonstrate the potential of graphene separation media, implementation as a viable membrane is limited by fabrication challenges. The limited membrane area produced through graphene synthesis^12^ combined with defects occurring during the transfer/assembly process^13,14^ present significant scalability challenges. GO laminate membranes prepared through vacuum filtration^15,16^ have been pursued as scalable alternatives. This process produces uniquely selective membranes due to a precise interlayer spacing between the GO nanoplatelets. However, GO membranes prepared through this method are generally thick, severely increasing the species transport path length (i.e. transport resistance) compared to a single-layer graphene membrane.

Layer-by-layer (L-b-L) assembly of GO^17,18^ has been pursued in an attempt to reduce the membrane thickness and transport resistance, while maintaining an acceptable selectivity. L-b-L assembly frameworks have achieved minimum thicknesses of ~100 nm. These assemblies have demonstrated water permeabilities of ~25-60 mL/hr.mmHg.m^2^ with improved retention of charged molecules compared to typical nanofiltration (NF) membranes^19,20^. However, L-b-L assemblies have largely ignored the impact of critical GO nanoplatelet physicochemical properties on the membrane transport characteristics. For example, as depicted in Fig. 1, the nanoplatelet size dictates the overall transport path length of the species through the membrane - large nanoplatelets reduce the permeability.

**Fig. 1:**
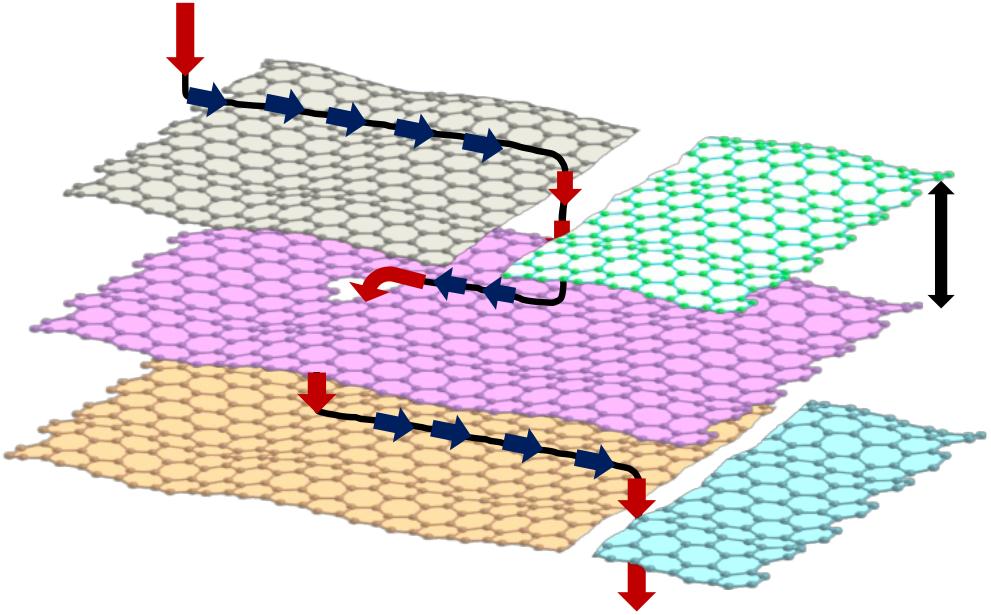
Illustration of species transport across GO laminate through nanoplatelet-edge and defects. Primary transport pathways through self-assembled GO nanoplatelet layers, dependent on nanoplatelet size and interlayer spacing.

An ideal GO membrane should consist of the fewest number of GO layers while avoiding defects that diminish the membrane selectivity. Vacuum filtration and current L-b-L assembly yield a wide array of nanoplatelet assemblies, ranging from sparsely packed (Fig. 2a) to overly condensed layers hardly exhibiting any order and planarity (Fig. 2b) necessary for minimizing through-plane defects, and consequently the number of GO layers. We have discovered that if small GO nanoplatelets (~100s nm) are given sufficient diffusion time to assemble, rather than the common dip-and-rise step used in the L-b-L assembly process, they can produce a closely packed, ordered GO mosaic (Fig. 2c) enabling a highly selective membrane with only a few GO layers.

**Fig. 2:**
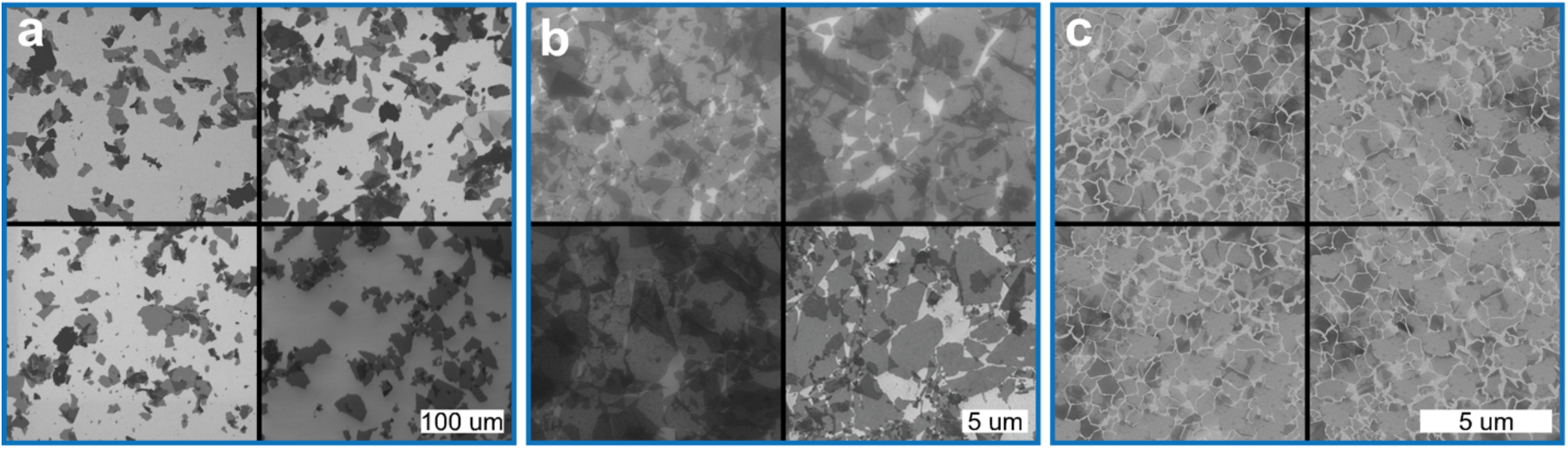
Visualization of GO nanoplatelet assembly utilizing various layering techniques. Nanoplatelet coverage and alignment using (**a &b)** conventional dipping processes or **(c)** layer-by-layer assembly. **a,** Sparse distribution of GO nanoplatelets after under-compression dipping. **b,** Over-compacted GO nanoplatelets through dipping. **c,** Ordered assembly of GO nanoplatelets through optimal layer-by-layer processing.

This approach has been inspired by our studies of clay nanoplatelets self-assembly^21^. The GO nanoplatelet organized self-assembly is dictated by the electrostatic interactions between the nanoplatelets, particularly their edge functional groups, and the interlinking molecules charged end group. In a solution with suitable pH, GO nanoplatelets tend to avoid overlapping due to electrostatic repulsion of their negatively-charged functionalities, while being attracted by the interlinker. The net result of this interaction is edge-to-edge and surface-to-surface repulsion between the GO nanoplatelets, and surface-to-surface attraction between GO nanoplatelets and the underlying substrate functionalized with charged molecules. In this case, a positive interlinker is selected to attract negative functionalities of the GO nanoplatelets. This process fundamentally differs from GO laminates prepared through vacuum filtration and dip-and-rise L-b-L assembly, where GO nanoplatelet orientation is wholly random due to mechanical forces overcoming the relatively weaker functional group repulsive forces. Providing nanoplatelets with sufficient time allows them to settle/orient and assemble. The non-interlinked GO nanoplatelets remained on the assembled/locked platelets while the sample is pulled out of the solution are then removed through submerging in a clean DI water bath followed by a gentle DI water rinse. This planar GO layer formed is subsequently functionalized with another layer of interlinking molecule, for assembly of the next GO layer.

The nanoplatelet size is often not reported in studies that have utilized L-b-L assembly. Limited studies have reported a size on the order of micrometers^22–25^. The Hummers’ and Marcano’s methods that are universally used to exfoliate GO produce nanoplatelets ranging from a few micrometers to 10s μm^26^. By breaking the GO nanoplatelets (Fig. 3a-c) through sonication, they can better diffuse and rotate/orient while forming smaller open areas (i.e. defects) in between themselves. Combining this highly ordered nanoplatelet assembly with an appropriate interlinking molecule (polyallylamine hydrochloride, PAH) enable highly permeable membranes with a precise molecular weight cut-off (MWCO). Fig. 3d provides an atomic force microscope (AFM) image of the GO mosaic showing a precise 5 nm distance from the underlying substrate.

**Fig. 3:**
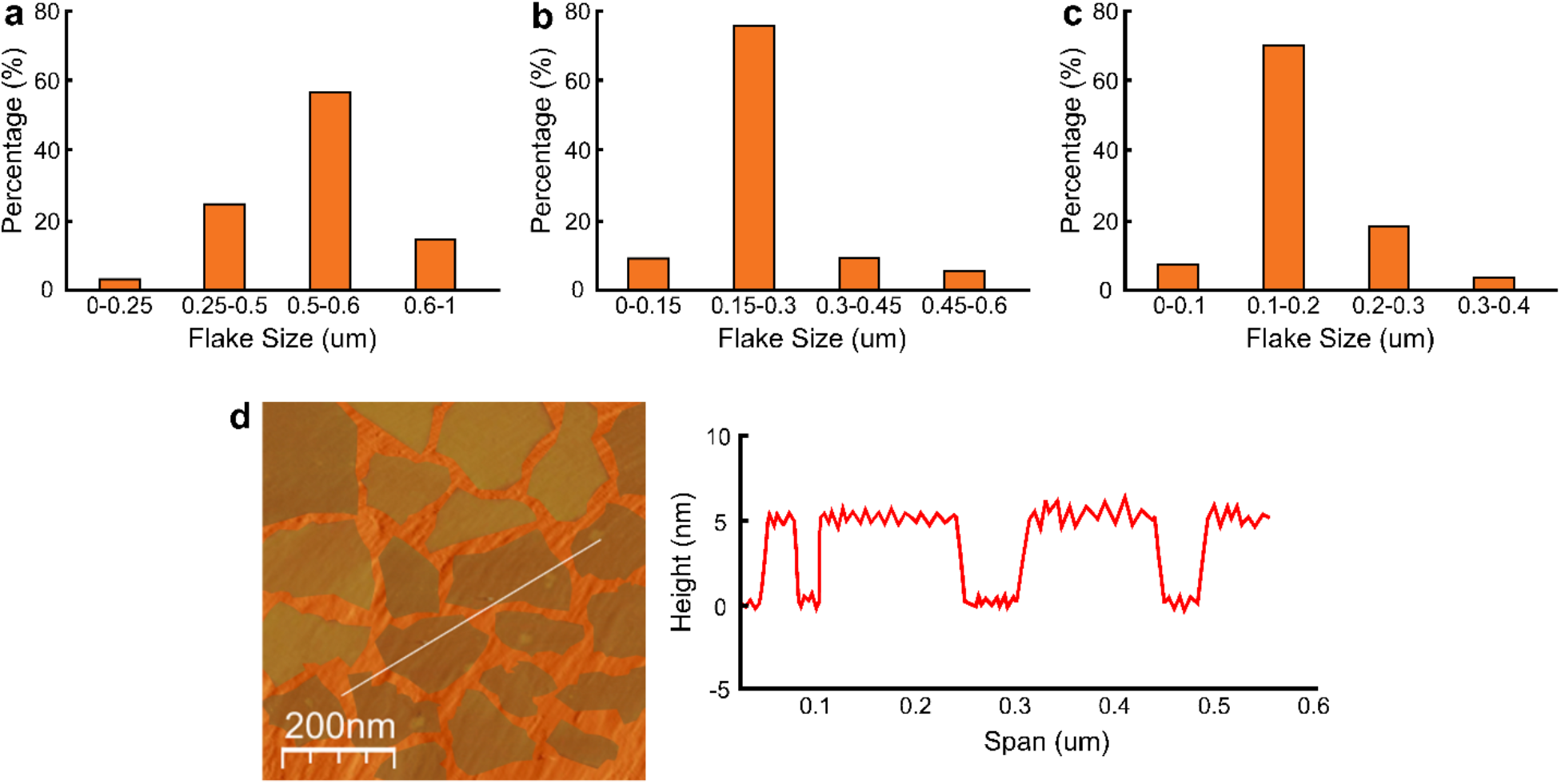
GO nanoplatelet size distribution and PAH interlinker characterization across tested devices. Nanoplatelet size distribution for **(a)** 500 nm support, **(b)** 250 nm support, and **(c)** 100 nm support. **d,** AFM characterization of PAH interlayer spacing for GO anchored on PMMA support.

An ideal assembly fabricated using the aforementioned process would only require two layers of GO, but this has yet to be achieved without incurring defects that compromise permselectivity. Herein, we demonstrate the efficacy of a membrane utilizing three GO layers. To achieve a high selectivity with only three GO layers, it was critical to use a near-atomically smooth underlaying substrate. Polyethersulfone (PES) membranes used in prior studies^27,28^ are very rough (see Supplementary Fig. 1), requiring lamination of layers of large GO nanoplatelets to establish planarity. Furthermore, the pore size of the support membrane must be smaller than the GO nanoplatelets and uniform to prevent nanoplatelets clogging of the pores. PES membranes have a wide range of pore size (Supplementary Figs. 1 and 2). Hence, three polymethyl methacrylate (PMMA) membranes with pore sizes of approximately 100, 200, and 400 nm (insets in Fig. 4a) were nanoimprinted and then hydrolyzed for the GO nanoplatelet sizes presented in Figs. 3a-c. Fig. 4b shows that the 1^st^ GO assembled layer provides significant coverage of the underlying PMMA membrane, but has large defects left between the nanoplatelets. The 2^nd^ layer (Fig. 4c) substantially reduced the defects but a 3^rd^ layer was needed to eliminate visible defects (Fig. 4d).

**Fig. 4:**
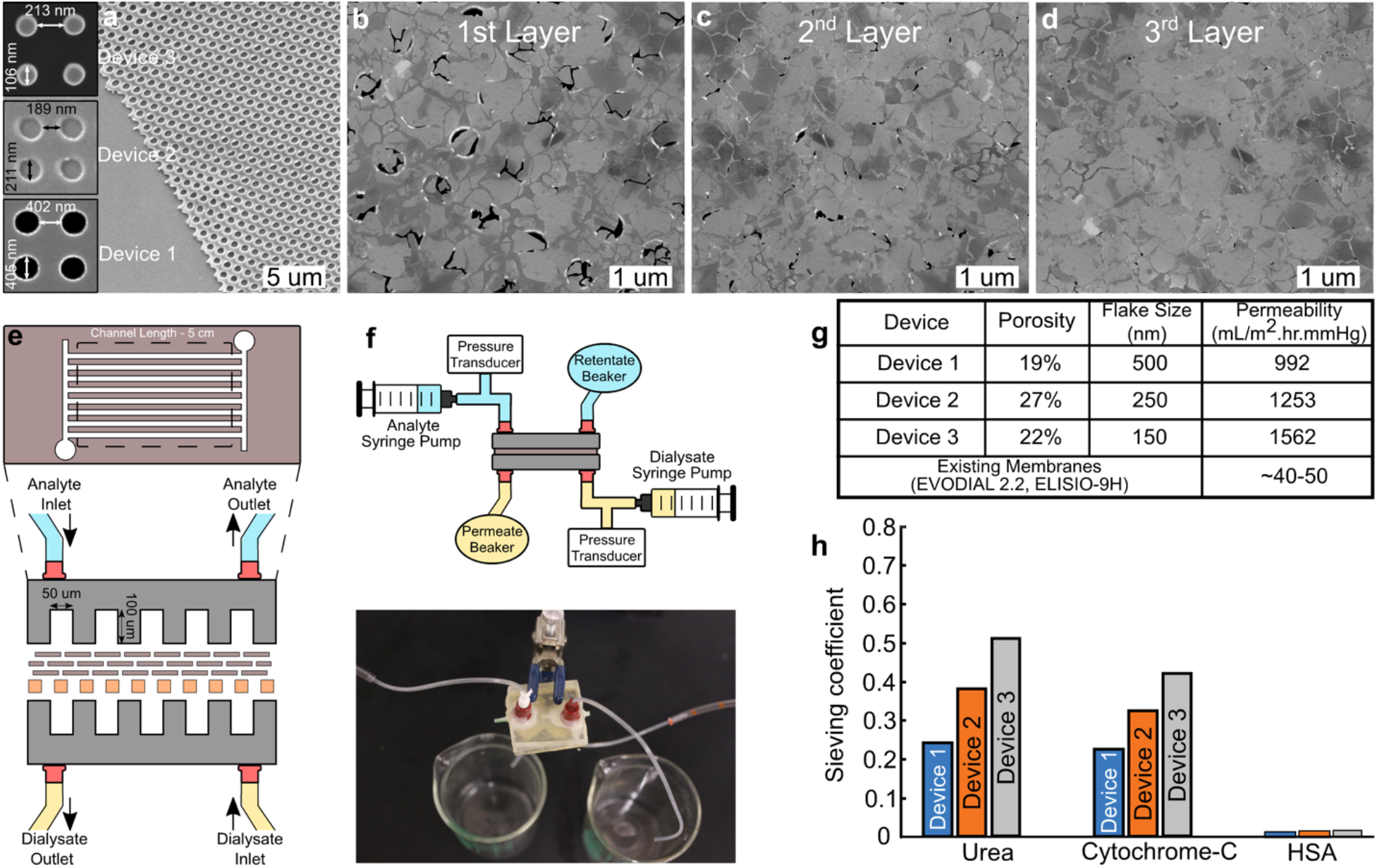
Visualization of PMMA support and impact of subsequent GO layering, with permeability and permselectivity results for tested device iterations. Porous PMMA support and subsequent GO bilayers assembled atop support. **a,** Schematic representation of the characterized GO device. **b,** System schematic for sieving characterization and actual device. **c,** Device iteration characteristics and permeability compared to typical commercial membranes. **d,** Preliminary sieving performance of low and middle-weight uremic toxins with minimal albumin loss.

The microfluidic device shown in Fig. 4e was used to determine the membrane transport properties. The water permeability data are provided in Fig. 4g. Reducing the nanoplatelets size directly improved the permeability, through shortening the effective transport path length, and a permeability of 1562 mL/hr.mmHg.m^2^ was measured with Device 3, two orders of magnitude higher than existing nanofiltration membranes. Such a membrane offers major value to the hemodialysis industry. The permeability of this membrane is nearly forty-fold higher than the commercial high flux hemodialysis membranes (e.g. EVODIAL 2.2 and ELISIO-9H manufactured by Baxter Healthcare Ltd. and Nipro Medical Corp., respectively). This permeability is nearly 5× greater than that of the glomerular membrane of the kidney^29^, demonstrating the unique capability of nanomaterials to exceed their biological equivalent. Another unique transport characteristic of the membrane is its precise MWCO that offers ultimate selectivity relative to polymer membranes that have a range of pore sizes.

The advantages over the existing technology of this membrane far outweigh its higher cost relative to the polymer membranes. With such a high permeability, only a fraction of a square meter is required for dialysis of a human adult, which can be built into a small multilayer microfluidic cartridge to reduce the extracorporeal blood circuit by an order of magnitude, preventing loss of 100-150 mL blood in each dialysis session. The new membrane module can be operated with hemodynamic pressure rather than an external pump that is a source of hemolysis due to the high internal shear forces. All these factors can enhance quality of care while reducing costs, particularly through increasing home dialysis; in-clinic care costs the Medicare system an average of $91,000 per patient^30^, with negligible contribution from the membrane module itself. This must be contrasted with other potential applications of graphene-based membranes such as desalination where tens of thousands of square meters^6^ membrane is required with the promise being reduced cost, not energy consumption that is dictated by the pressure required to counter the osmotic pressure.

The membrane sieving capability of urea and cytochrome-c as representative small and middle-weight uremic toxins, while monitoring the retention of albumin, are provided in Fig. 4h. Permeate and sieving characteristics are assessed using a custom setup. The GO test device was a 5-cm-long single serpentine polydimethylsiloxane (PDMS) microchannel with a 50 × 100 μm^2^ cross-section. The device was connected to two syringe pumps (Fig. 4f) used to deliver the analyte and dialysate at the same flow rate of 1 mL/hr (i.e. 0.017 mL/min). PX-26 pressure transducers were utilized in-line to monitor the pressures within the system. Sieving performance of the device is assessed using urea (4.6 mmol/L), cytochrome-c (0.08 mmol/L), and human serum albumin (0.075 mmol/L). A maximum urea sieving coefficient of 0.5 is achieved for Device 3 while the human serum albumin (HSA), with a size of <66 kDa, retention was >99% across all devices (Fig 4h). Considering the surface area of the GO membrane (i.e. 2.5 mm^2^) used in our test device, 0.015 m^2^ effective sieving area is necessary for nocturnal dialysis with a flow rate on the order of 100 mL/min. This membrane area can be incorporated in a microfluidic membrane module with a 5 × 5 cm^2^ footprint consisting of 15 microchannel layers.

Next, we investigated the hemocompatibility of GO-based membranes compared to commercially available hemodialyzer materials. The GO-blood interactions are investigated, utilizing careful control on oxidation extent and GO nanoplatelet size to identify their hemocompatibility. GO was synthesized^31,32^, sonicated in a bath sonicator^33,34^, and then deposited onto glass substrates, where complete coverage of the glass substrate with GO nanoplatelets is critical in preventing the substrate contribution toward hemolysis. SEM imaging shows close packing of GO nanoplatelets on the glass substrate with minimal exposure of the underlying glass, where layers of GO-PAH fully cover the surface.

First, we investigated hemolysis, the rupture of red blood cells, following 1 hour of exposure to membrane surfaces. We found that diethylaminoethyl cellulose (DEAE) and regenerated cellulose (RC) membranes, which fell out of favor as hemodialysis materials, induced a slight level of hemolysis (~2%), as expected. For these tests, 1 mL of the hematocrit-adjusted blood (*Hct* = 36%) was pipetted onto each sample. With 1 mL of blood, the total surface area to blood volume ratio is approximately 5.4 cm^2^/mL, following the specifications recommended by ASTM F756^35^. The 1 mL blood, when spread over the 5 × 1 cm^2^ area, did not confer sufficient hydrostatic pressure to drive the seepage of blood through DEAE, RC, or PES membranes, enabling straightforward recovery for analysis similarly to the GO coated test substrates. These membranes produced notably higher hemolysis compared to all four tested GO oxidation assemblies (Fig. 5a). Teflon, silicone, and glass, which were used in the GO testing apparatus, showed comparable hemolysis levels compared to the different GO substrates indicating that the actual GO contribution to hemolysis may be even lower than that measured.

**Fig. 5:**
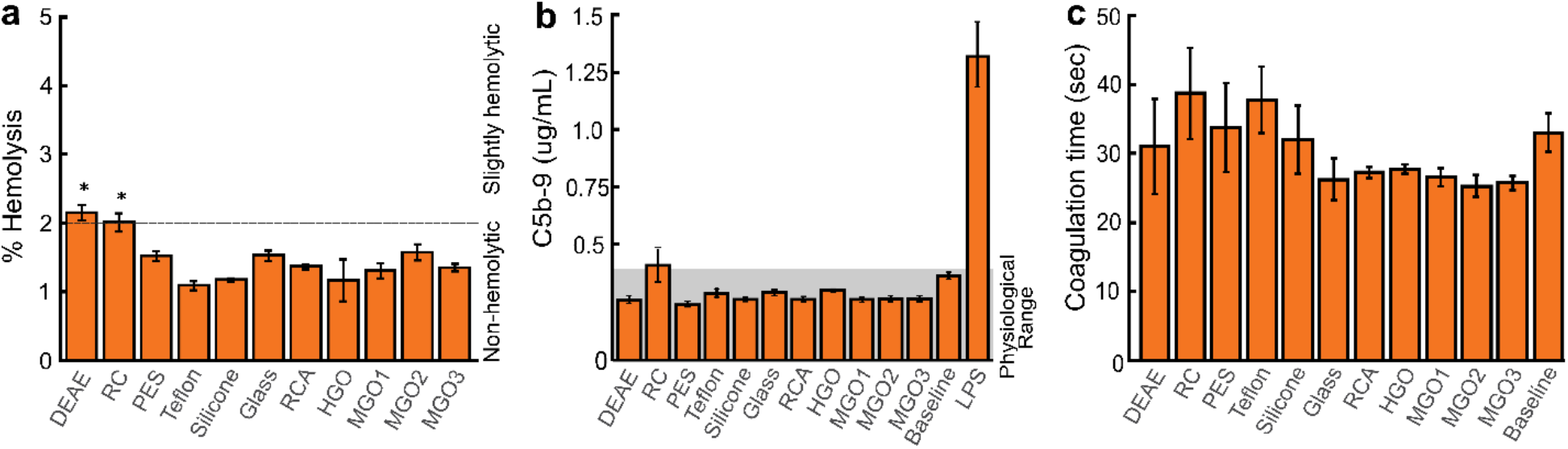
Hemolytic and C5b-9 complement activity for GO suspension and bilayer scenarios. **a,** Hemolysis results for GO membrane conformation with varying oxidation factors and comparative commercial substrates. **b,** Complement activation results for GO membrane layout (Each bar and the associated error bar represent the mean and the standard error of six independent samples (n = 6)). For positive control (LPS) and the negative controls at 4°C and 37°C, n = 3. **c,** Coagulation results for GO and standard membrane material.

We then investigated substrate-induced immunogenicity, as assessed by the production of C5b-9 complement using a similar frame test structure to the hemolysis studies. The test samples were incubated for 15 minutes in a manner similar to that described in the hemolysis testing. 10 μL of the plasma sample was retrieved and diluted 1:30 in PBS. 100 μL of the diluted sample was then assessed for complement activation using a C5b-9 ELISA (enzyme-linked immunosorbent assay) kit. C5b-9 ELISA revealed (Fig. 5b) that none of the test substrates promoted significant C5b-9 production, with all GO samples falling within the physiological range. We induced complement activation using 100 ng/mL of lipopolysaccharide (LPS) from E. coli O55: B for a positive control, which resulted in markedly increased levels of C5b-9 production. There was no statistical significance between GO substrates and PES membranes.

Thrombogenicity, or the tendency of a material to induce clotting, of the GO surface was evaluated based on coagulation time after post-thrombin addition, where shorter times for coagulation onset corresponded with higher thrombogenicity. No statistically significant differences were observed across all GO variants (Fig. 5c) compared to the control substances. Compared to polyethersulfone (PES) membranes and Teflon, noted for their highly hemocompatible characteristics, GO membranes exhibit no significant variance in hemolytic, coagulation, or complement activation characteristics (Fig. 5a-c).

These hemocompatibility results are contradictory to prior GO compatibility studies^36,37^, which indicated that GO induces high levels of hemolysis and complement activation^38–40^. Comparison between GO suspension and membrane platforms suggest that the interactions between red blood cells (RBCs) and GO platelets in these two cases fundamentally differ. The ability of GO nanoplatelets in suspension to freely diffuse leads to an increased interaction rate with other species. A nanoplatelet distribution ranging from 150-500 nm was analyzed through nanoparticle tracking analysis (NTA). When freely diffusing, a randomly oriented GO nanoplatelet with an estimated disc diameter of 150 nm has a diffusion coefficient of ~2.9 μm^2^/sec. At an RBC concentration of 5 × 10^8^/mL and a GO concentration of 3.5 × 10^10^/mL, this translates to a GO-RBC encounter frequency of ~82 times every 20 sec. This interaction frequency is drastically higher compared to the bilayer scenario that we present in this study, as RBC sedimentation tends to occur, limiting the number of RBCs which can actively interact with the surface. Even when accounting for recirculation of the RBCs atop a GO laminate at 10 dyn/cm^2^ that might occur in a wearable hemodialyzer, the hemolytic activity is comparable to polymer baselines and well below GO suspensions (Fig. 6c). These results suggest that the primary hemolytic mechanism found in suspension studies is absent in GO laminates or occurs on a much less pronounced magnitude.

**Fig. 6:**
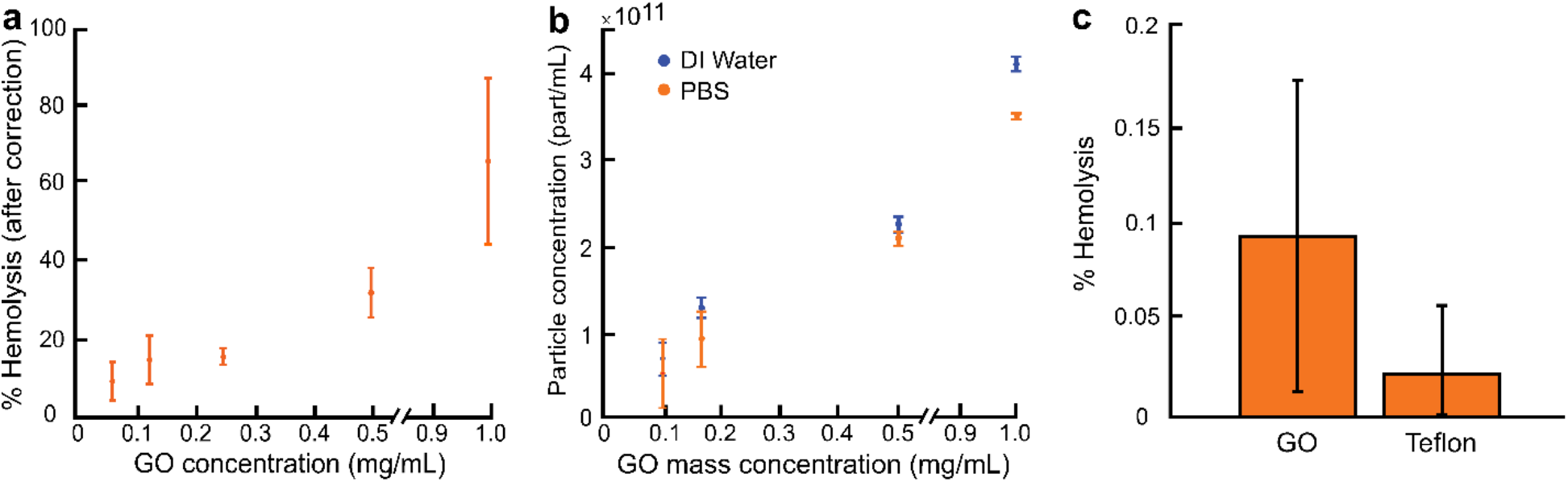
Characterization of GO suspension behavior and quantification of hemolytic behavior after perfusion at physiological conditions. **a,** GO suspension hemolysis using GO 60-minute sonication at varied concentrations. **b,** GO aggregation in DI Water and PBS solutions based on number of particles present with higher aggregation in PBS. **c,** Hemolysis observed after perfusion across GO surfaces, which falls in the non-hemolytic regime (<2%).

In summary, we demonstrated a unique self-assembled GO nanoplatelet ordered mosaic, greatly advancing a decade-old effort on development of graphene-based membranes. Careful control of GO nanoplatelet characteristics and the self-assembly process along with the use of a suitable planar and smooth support layer were key to this accomplishment. The new membrane requires only three layers of GO atop a PMMA support, achieving permeabilities as high as 1562 ± 30 mL/m^2^.hr.mmHg, nearly two orders of magnitude greater than existing nanofiltration membranes. A precise effective pore size of 5 nm, a unique attribute of GO laminates considered very significant in separation domain, represents a great advantage over the polymer membranes with a range of pore sizes. This GO laminate has also shown vastly improved hemolytic and biocompatible properties compared to previous studies concerning GO nanoplatelets in suspension. Even under recirculation conditions of 10 dyn/cm^2^, hemolytic activity of GO laminates remains at or below the commercially available dialyzers. The membrane provides a viable platform for miniaturized dialysis devices that could enhance in-home low flow rate nocturnal dialysis.

## Methods

### Substrate-induced hemolysis

The GO and control substrates were made to match the format of a glass side (7.5 cm × 2.5 cm). Due to the manufacturing process involved in the preparation of GO, the front and back sides of GO samples were not always identical. We created a custom-cut framework (300 μm-thick, 5.8 cm × 1.8 cm in area, with 5 cm × 1 cm cutout in the middle) using restricted grade medical silicone to hold whole blood on top of the samples for incubation. The surface area of the silicone gasket (from the 4 walls) is ~7% of the total surface area that will be exposed to blood. For control substrates such as the DEAE, RC, and PES membranes, 5 cm × 1 cm cutouts were placed on top of Teflon (7.5 cm × 2.5 cm) and held in place within the silicone frame.

Each sample was hosted within a petri dish to better ensure sterility and kept in an incubator (37°C, 5% CO_2_, 80% relative humidity) for 2 hours. Each petri dish also contained a wetted Kimwipe to help reduce blood evaporation. We observed minimal seepage of blood onto Teflon after the 2-hour incubation. The blood on top of the sample was gently mixed through titration and retrieved (700 μL) for hemoglobin (Hb) measurement. The 700 μL sample was centrifuged at 500 G for 20 minutes. The top 200 μL of the resulting plasma was then retrieved and centrifuged again at 3,000 G for 10 minutes to further ensure the removal of RBCs. The top 100 μL of the resulting plasma was placed into a 96 well plate for hemolysis assessment.

### Substrate-induced complement activation

For complement activation testing, the plasma was kept at 4 °C (on ice) at all times except during the sample incubation. We employed a similar strategy of holding the plasma (150 μL) on top of the sample using custom-cut frames made of restricted grade medical silicone (300 μm-thick, 2 cm × 6 cm in area, and contained 3 circular cutouts, each 1 cm in diameter). Since the kit did not provide the 96 well plates, we used the 96 well EIA/RIA plate from another vendor (CoStar, Washington DC). 1:30 dilution was performed to bring the C5b-9 level down to the detection range of the ELISA kit. The sample incubation in the ELISA plate was performed on an icepack to reduce unintended complement activation by the ELISA plate itself. Sample incubation time on the ELISA plate was also shortened from 2 hours to 15 minutes. Otherwise, the ELISA is carried out following manufacturer instruction.

### Substrate-induced coagulation

We assessed coagulation based on thrombin time (TT). Thrombin (0.5 Unit/mL) was added to fresh platelet poor plasma (PPP) to initiate the coagulation cascade and coagulation time was recorded. We used the setup of the platelet aggregometer (CHRONO-LOG Model 700) to conduct the coagulation testing. In this setup, the plasma was maintained at 37 °C in a narrow glass cuvette and well mixed with a siliconized magnetic stir bar. The time at which the stir bar stopped moving was recorded as the coagulation time.

### Statistical analysis

Hemolysis and GO nanoplatelet studies were carried out in three independent studies. Complement activation samples were evaluated in six independent studies.

## Data Availability

The data that supports the findings of this study are available from the article and Supplementary Information, or from the corresponding author upon request.

## Acknowledgements

Research reported in this paper was supported by NIBIB of the National Institutes of Health under award number R21EB023527 to S.M. and T.R.G. The content is solely the responsibility of the authors and does not necessarily represent the official views of the National Institutes of Health. We thank Bradley Kwarta for creating the GO laminate illustration.

## Author Contributions

S.M. and T.R.G. were responsible for the project conception. R.P.R. was responsible for all GO-based preparations and H.H.C. performed biocompatibility experiments. H.N.M performed NTA analysis experiments and assisted with hemocompatibility experiments. R.P.R and H.H.C wrote the manuscript and supplemental information with contributions from all authors.

## Additional Information

Reprints and permissions information is available at www.nature.com/reprints. The authors claim no competing interests. Correspondence and requests for materials should be addressed to S.M. or T.R.G.

## Supplementary Information

**Supplementary Fig. 1.**
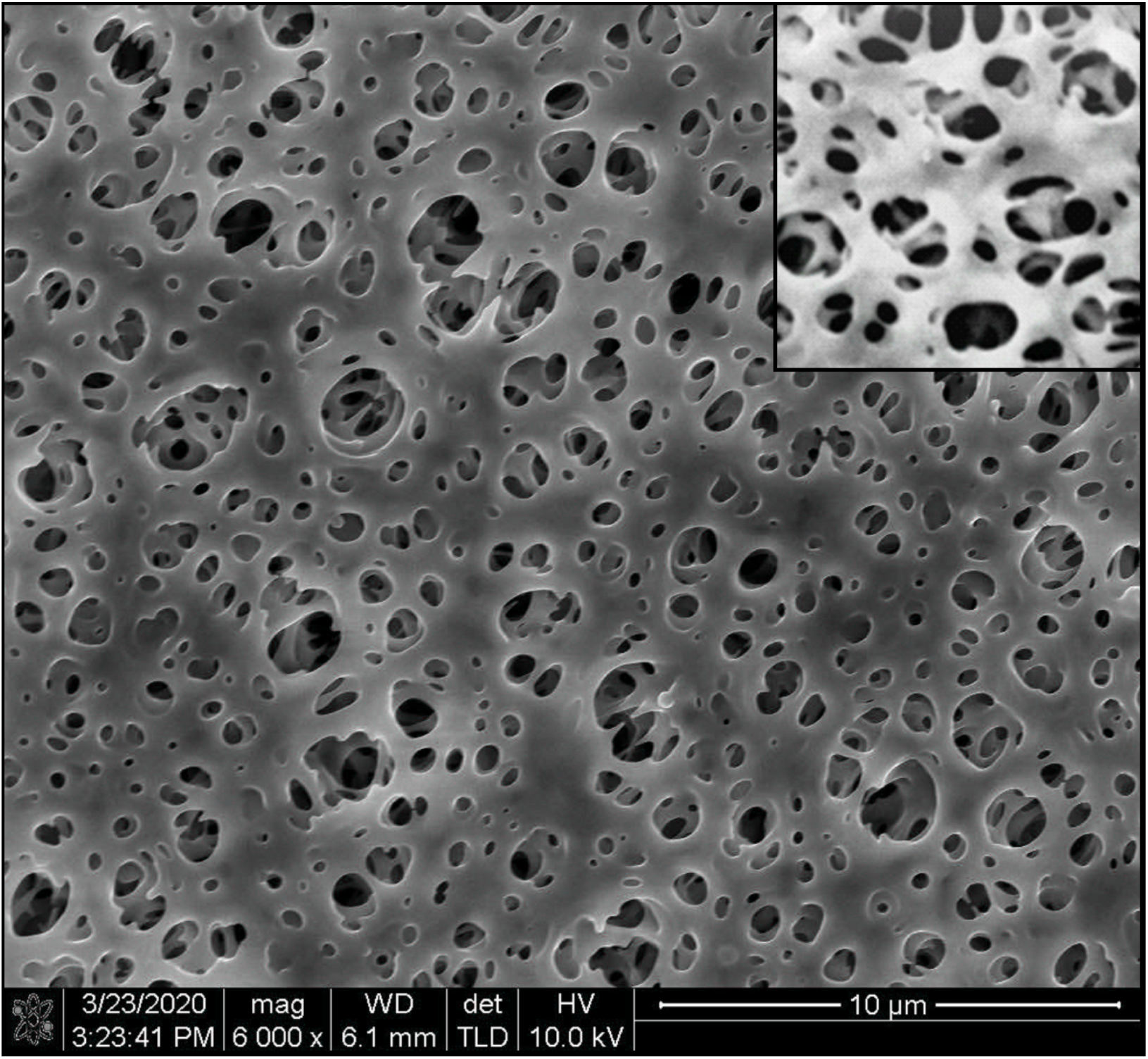
Surface roughness and porosity visualization of PES membrane. SEM imaging of PES surface structure/porosity with magnified inset image showing variation in surface smoothness.

**Supplementary Fig. 2.**
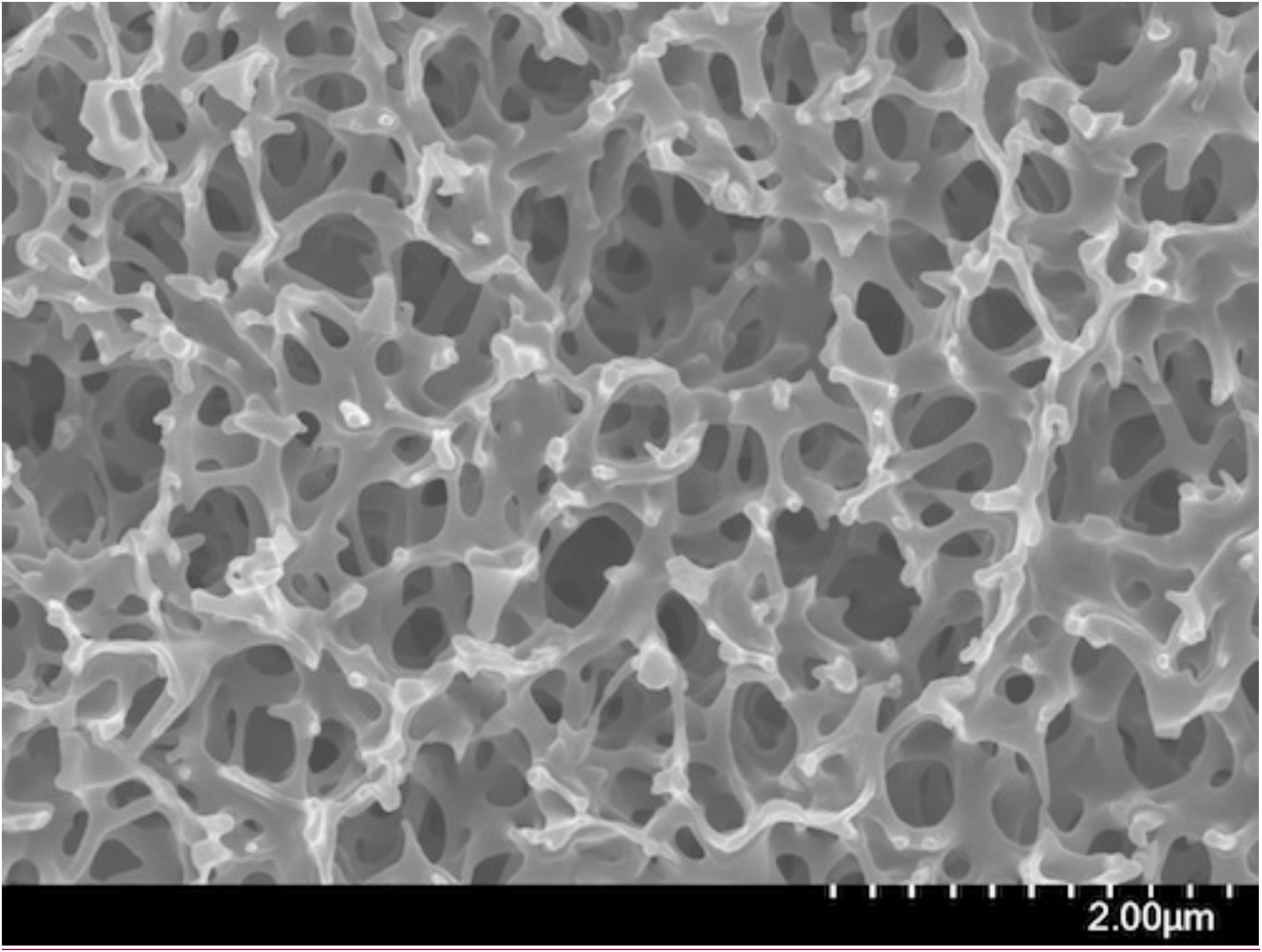
Roughness imaging of PES membrane. High-magnification imaging of PES surface.

## Methods

### Langmuir-Blodgett nanoplatelet analysis

Graphene oxide (GO) nanoplatelet size was characterized using typical Langmuir-Blodgett deposition^1^ onto a silicon wafer imaged via SEM. ImageJ software was used to quantify nanoplatelet size distribution.

### Fabrication of microchannel test device

~ 200 nm thick PMMA support was spin-coated onto a silicon wafer and subsequently imprinted across three separate devices to introduce 400, 200, and 100 nm pores. The porous PMMA support was then bonded to a PDMS substrate containing microchannels (50 μm wide x 100 μm deep) over 5 cm length for fluid delivery. This PDMS substrate was bonded through oxygen plasma treatment to the porous PMMA to seal the bottom portion of the device. This PDMS-PMMA structure was removed from the underlying silicon wafer through etching of the aluminum sacrificial layer. PAH was used as the cationic interlinking molecule, where the hydrolyzed PMMA support was immersed in PAH or GO solution (1 g/L) for 15 minutes to introduce subsequent layering. After successful assembly of three PAH-GO bilayers atop the PMMA support, an upper PDMS microchannel was epoxy bonded to the lower device to complete the device fabrication.

Once the GO device was fully fabricated, permeate and sieving characteristics were assessed using an in-house setup. Pressure transducer data was monitored by data acquisition system (Agilent Technologies Inc., CA) at room temperature (T = 20 - 25°C). 0.5 mL samples were collected for analysis from the analyte and permeate outlets every 30 minutes. Urea sample concentrations were assessed using urea colorimetric assay kit (Biovision Inc., CA) combined with micro-plate spectrophotometry. Cytochrome-c and albumin concentrations were determined through UV-Vis spectroscopy at 405 nm and 280 nm, respectively.

### Fabrication of microfluidic device for hemocompatibility evaluation

A simple microfluidic device was created by bonding a PMMA cap to the substrate (GO or Teflon) using double-sided adhesive transfer tape (3M 468MP, by 3M, St. Paul, MN). The PMMA cap was cut from 0.56 cm thick PMMA sheets using a 90W CO2 laser cutter (Full Spectrum P48-36, by Full Spectrum, Las Vegas, NV) at the RIT Maker Space. The PMMA top piece contained 2.85 mm holes to host the insertion of E-3603 tygon tubing (ID/OD = 1/16”/3mm, by Saint-Gobain, Malvern, PA), which enabled fluidic access to the substrate. Rectangles (2 mm × 5 cm) were cut from the adhesive transfer tape (~110-130 μm thick) using a digital craft cutter (Silhouette Cameo, by Silhouette America, Lindon, UT). Four microfluidic channels were situated across the 2.5 cm × 7.5 cm substrate footprint.

### Blood storage and handling

Blood anticoagulated with 3.2% (v/v) sodium citrate were purchased from ZenBio (Research Triangle Park, NC) and immediately aliquoted in 50 mL polypropylene centrifuge tubes (Becton, Dickinson and Company, Franklyn Lakes, NJ), stored at 4°C, and used within 21 days. For hemolysis testing, the aliquoted blood (40 mL) was warmed to room temperature for at least 30 minutes and diluted with PBS to adjust the hematocrit (Hct) to 36%. Plasma was used in place of blood for complement activation testing. The plasma was obtained as supernatant after the aliquoted blood was centrifuged at 1200 G for 10 minutes at 4°C. In the case of coagulation testing, platelet rich plasma (PRP) was used instead of whole blood. Fresh whole blood was collected in blue top vacutainers (Becton, Dickinson and Company, Franklyn Lakes, NJ), which contained the anticoagulant sodium citrate, and kept at room temperature. The PRP was obtained as supernatant after the fresh whole blood was centrifuged at 150 G for 15 minutes.

### Substrates

The control substrates used for the hemocompatibility study, along with their manufacturer product number, were provided as follows: glass (VWR Cat No. 48300-025); DEAE: DE81 grade chromatography Diethylaminoethyl cellulose (Whatman® Cat No. 3658-325); RC: 100 kDa Regenerated cellulose (Ultracel® Cat No. PLHK02510); PES: 100 kDa ultrafiltration polyethersulfone (Biomax® Cat No. PBMK02510); Teflon: “FAT” Teflon from CS Hyde (Cat No. 15-20A-1.5-5); silicone: restricted grade medical silicone (Specialty Silicone Fabricators, Paso Robles, CA).

### Hemolysis induced by GO in aqueous suspension

The protocol for measuring GO-induced hemolysis was modified from the work of Monasterio et al.^2^ Briefly, red blood cells (RBCs) were washed and re-suspended in Ca^2+^-free HBSS buffer in a stock concentration of 5 × 10^8^ RBC/mL. GO (6 mg/mL) and RBC stock were first mixed 1:6 to bring the GO concentration down to 1 mg/mL. This solution was then serially diluted 1:2 in RBC stock to vary the GO:RBC ratio. After 1 hour in the incubator (37°C, 5% CO_2_, 80% relative humidity), the GO/RBC mixtures were centrifuged at 12,000 G for 10 minutes. The top 100 μL of the supernatant were diluted 1:10 in deionized water (diH_2_O) prior to the Hb measurement. We used the Cripp’s method^3^ of Hb determination, which describes the relation between Hb concentration (in mg/dL) in terms of absorbance at 576.5 nm, 560 nm, and 593 nm in accordance to the following equation:

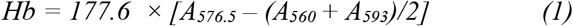

Ideally, the concentration of Hb released into the supernatant is proportional to the number of RBC that lysed. The degree of hemolysis is often reported in terms of a percentage, in accordance to the following equation:

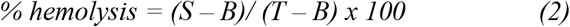

where *S* = Hb of the sample, *T* = Hb of lysed sample, and *B* = Hb measured at the beginning of the experiment.

*T* is obtained by intentionally lysing the whole blood, which should release all Hb from RBCs. Complete lysis was achieved either with 1:5 dilution of whole blood in diH_2_O or with the addition of 1% (v/v) Triton X. It was noted by Monasterio et al. that GO does bind away Hb at the physiological pH (7.4). Therefore, after centrifugation, the pelleted GO will sequester away some (if not all) of the Hb to produce a low apparent level of hemolysis. To address this artifact, a correction curve was obtained by incubating known concentrations of GO with known concentrations of Hb for 1 hour to assess the Hb binding capacity of GO. We also noted that the initial 1:6 dilution of the GO in RBC stock potentially produces an artificial background lysis since GO was made in diH_2_O suspension. To address this concern, we created a vehicle control with 1:6 dilution of diH_2_O in RBC stock and found minimal level of hemolysis (< 0.1%).

### Hemolysis induced by RBC perfusion over the substrates

RBC stock (5 × 10^8^ RBC/mL in HBSS) was perfused through the microfluidic device at a flow rate of 250 μL/min. At room temperature, this produced a wall shear stress of ~10 dyn/cm^2^ near the substrate surface. The RBC stock exiting the microfluidic device was collected every 8 minutes (2 mL per collection) and assessed for hemolysis.

### Perfusion-induced GO delamination

Ultrapure water was perfused through the microfluidic device at varying shear stress. The number and the size of the particles in the exiting water were determined using nanoparticle tracking analysis (NanoSight NS300, by Malvern Panalytical Ltd, Malvwern, UK).

